# Pre-existing visual responses in a projection-defined dopamine population explain individual learning trajectories

**DOI:** 10.1101/2024.02.26.582199

**Authors:** A Pan-Vazquez, Y Sanchez Araujo, B McMannon, M Louka, A Bandi, L Haetzel, International Brain Laboratory, JW Pillow, ND Daw, IB Witten

## Abstract

Learning a new task is challenging because the world is high dimensional, with only a subset of features being reward-relevant. What neural mechanisms contribute to initial task acquisition, and why do some individuals learn a new task much more quickly than others? To address these questions, we recorded longitudinally from dopamine (DA) axon terminals in mice learning a visual task. Across striatum, DA responses tracked idiosyncratic and side-specific learning trajectories. However, even before any rewards were delivered, contralateral-side-specific visual responses were present in DA terminals only in the dorsomedial striatum (DMS). These pre-existing responses predicted the extent of learning for contralateral stimuli. Moreover, activation of these terminals improved contralateral performance. Thus, the initial conditions of a projection-specific and feature-specific DA signal help explain individual learning trajectories. More broadly, this work implies that functional heterogeneity across DA projections serves to bias target regions towards learning about different subsets of task features, providing a mechanism to address the dimensionality of the initial task learning problem.

## Introduction

A key challenge of learning a new task is that the environment is high dimensional - there are many different sensory features and possible actions, with typically only a small subset reward-relevant. While animals can learn to perform complex tasks that involve arbitrary associations between stimuli, actions and rewards^1–6^, a consistent and striking result across varied experimental paradigms is that in initially acquiring such tasks, large differences between individuals are apparent in the learning process^7–12^. For example, although genetically inbred mice achieve high and consistent steady-state performance in a standardized visual decision-making task^7^, there are vast differences across individuals in how many training sessions are required to reach this state. What neural mechanisms contribute to initial task acquisition, and why do some individuals learn a new task much more quickly than others?

Midbrain dopamine (DA) neurons that project to the striatum have been implicated in learning^13–15^ and in individual differences^16–22^. However, DA is often studied in animals that are already familiar with a task’s structure; thus, whether and how DA explains individual differences in *de novo* task acquisition is not clear.

Given that DA neurons encode reward predictive cues^15^, one might expect task-related DA responses and behavioral sensitivity to those task events to co-evolve during task acquisition, as animals learn that task events are reward predictive^9,23–26^. This would be consistent with DA reinforcement signals reflecting (and possibly even supporting) task acquisition.

However, such a result would not in itself explain *why* different individuals progress along different learning trajectories. Instead, individual differences in the DA system that *precede* learning differences would point to a mechanism with the potential to explain differences in learning trajectories.

In considering the contribution of DA to individual differences in task acquisition, it may be informative to consider differences across projection-defined DA populations^16,17,20,21,27^. While traditionally the midbrain DA system was considered to provide a global, homogeneous reinforcement signal to the striatum^15^, there is increased appreciation of functional heterogeneity across dopamine neurons, with at least some of this heterogeneity varying by striatal target^27–35^. Such heterogeneity may in part reflect the high dimensionality of stimuli and actions^36,37^, with different DA subpopulations specializing in subsets of potentially relevant features.

Here, we performed longitudinal fiber photometry from DA terminals across 3 striatal subregions as mice acquired *de novo a* standardized visual decision-making task^7^. We found that learning trajectories were idiosyncratic, with different individuals exhibiting different patterns of side-specific learning. Across the striatum, DA responses to the visual stimuli increased with learning and closely tracked side-specific learning trajectories, consistent with widespread reward prediction error coding. Strikingly, even before animals experienced any training, activity in DA terminals in the dorsomedial striatum (DMS) displayed visual-contrast-dependent responses to contralateral stimuli that predicted the extent of contralateral-side-specific learning. Moreover, optogenetic activation of DMS DA terminals at the onset of contralateral stimulus presentation improved contralateral performance. Thus, the initial conditions of a projection- and feature-specific DA signal help to explain individual differences in learning trajectories. More generally, this work implies that projection-specific heterogeneity in the DA system could serve to bias different striatal subregions towards learning associations with a subset of task features. This could simplify the problem of initial task acquisition by effectively reducing the dimensionality of the learning problem faced by each subcircuit.

## Results

### Idiosyncratic and side-specific learning of a standardized visual decision-making task

To study individual differences in learning, we leveraged a highly standardized visual decision-making task^7,38–40^. In this task, at the beginning of each trial, a visual grating was presented on a screen on either the right or left side. Mice received a reward by turning a steering wheel with their front paws in the direction that centered the grating on the screen (Fig. 1a, Supp. Fig. 1a-c). Correct wheel turns were rewarded with 3 μl of sucrose water, while incorrect ones lead to a short timeout (2 s) and white noise (0.5 s). Across trials, the visual gratings varied in contrast (100%, 25%, 12%, 6.25%), with contrasts randomly and uniformly selected. To maximize interpretability of the neural recordings over the course of learning, we did not use a shaping procedure^7^, a de-biasing protocol^7,41^, or any other experimenter intervention.

**Figure 1.**
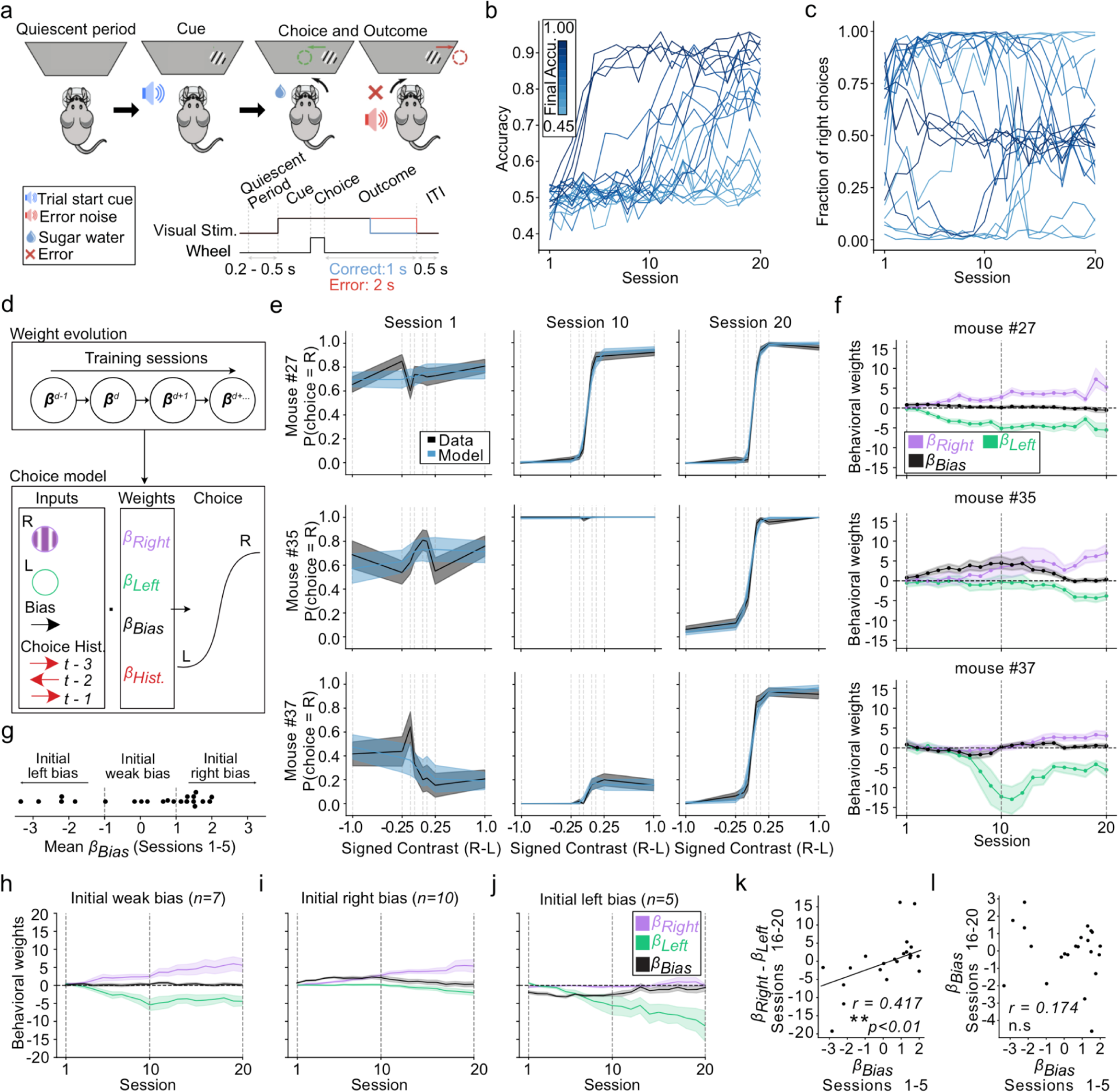
Idiosyncratic and side-specific learning trajectories. **a.** Schematic of the task. On each trial, a Gabor patch of a different contrast (6.12%, 12.5%, 25% or 100%) is presented on the right or left side of a screen. Centering the patch with a steering wheel leads to a small water reward while moving it out of the screen results in a short timeout (2 s) and white noise (0.5 s). **b.** Accuracy (fraction of rewarded trials per session) across training sessions. **c.** Probability of right choices across training sessions. In (b,c) each line represents one mouse, colored by their mean accuracy in sessions 15-20. **d.** Schematic of the behavioral model. Choice (left or right) on each trial is predicted with a logistic function based on weighting the contrast of the right and left visual stimulus (*β_right_ and β_left_*), a bias term (*β_bias_*) coded such that positive values indicate rightward choice, and a choice history kernel. Weights evolved across sessions (see Methods for details). **e.** Psychometric curves (“data”) and model predictions (“model”) from 3 example mice on the first, middle and last session of training. Lines and shading represent mean +/- s.e.m. **f.** Model weights across training for the same mice from (e). Lines and shading represent mean and 95% confidence intervals. **g.** Early *β_bias_* (average of sessions 1-5) for all the mice, showing the subdivisions used in subsequent panels between mice with weak, left, or right initial bias. **h-j.** Average trajectories of bias, right and left stimulus weights across training for mice subdivided by their initial bias as shown in (g). Lines and shading represent mean +/- s.e.m. across mice. **k.** Relationship between early *β_bias_* (average of sessions 1-5) and the late difference in stimulus sensitivity weights (*β_right_ - β_left_* for sessions 16-20). *r*=0.417, *p*=0.0007. **l**. Relationship between early *β_bias_* (sessions 1-5) and late *β_bias_* (sessions 15-20). *r*=0.174, *p*=0.522. In (k,l), each dot is a mouse. Correlation and p-values from robust regression. ** p <.01, ns: not significant. Across all panels, *n=22* mice.

Mice showed a large degree of variability in their learning trajectories, both in terms of the number of sessions to reach high accuracy (Fig. 1b) as well as in their probability of choosing left versus right wheel turns (Fig. 1c). While some mice selected both options (right and left) to an equal extent from the beginning of training, others appeared to prefer one side or the other.

To quantify choices across learning, we constructed a behavioral model (Fig. 1d) that explained each session’s contrast-dependent choices (i.e. psychometric curve) based on weights that evolved across sessions: a weight on the visual stimulus contrast on the left (*β_left_*) or right (*β_right,_*) that captured the contrast-dependent tendency to turn the wheel in the direction required by each stimulus, a choice bias (*β_bias_*) that captured the tendency to choose one side or the other irrespective of the stimulus, and a choice history regressor (*β_hist_*) that captured the tendency to repeat previous choices(see Methods for model details). This successfully reproduced the diverse psychometric curves observed across learning (Fig. 1e; 3 example sessions from 3 example mice) and allowed us to isolate the contribution of each variable to each mouse’s behavior across learning (Fig. 1f).

The model revealed that while some mice learned to increasingly weight visual stimuli similarly on both sides as training progressed (e.g., show an increase across training in both *β_right_* and *β_left_* in Fig. 1f, top), many instead displayed idiosyncratic learning trajectories where they preferentially weighted one stimulus versus the other (e.g. large *β_right_ vs β_left_* in Fig. 1f, middle; large *β_left_ vs β_right_* in Fig. 1f, bottom). This side-specific learning is consistent with the choice asymmetries evident in the raw data (Fig. 1c), though note that the model enables distinguishing between stimulus contrast-sensitivity (the stimulus weight) from a simple choice bias. The side-specificity of stimulus learning at the end of the training (*β_right_ - β_left_*) was predicted by the bias parameter *β_bias_* at the beginning of training, as evidenced both by plotting the stimulus weights based on groups defined by the level of initial bias (Fig. 1g-j) and by correlating initial bias with the difference between the final stimulus sensitivity weights (Fig. 1k). In contrast, early *β_bias_* did not predict final *β_bias_*(Fig. 1l).

Based on these observations, we concluded that mice display idiosyncratic side-specific learning trajectories that could be predicted by initial bias. We next explored how individual differences in striatal DA signals might explain these individual learning patterns.

### Across the striatum, contrast-dependent dopaminergic visual responses tracked side-specific individual learning trajectories

We recorded striatal DA signals longitudinally over the course of learning using fiber photometry. To ensure consistent expression of the activity indicator across time and individuals, we used double transgenic mice that expressed GCaMP6f in DA neurons (DAT-Cre x Ai148). Before training, each mouse was implanted with optical fibers into the following striatal subregions: dorsomedial striatum (DMS), dorsolateral striatum (DLS), and nucleus accumbens core (NAc; Fig. 2a, Supp. Fig. 2a-c). Each subregion was recorded unilaterally, with a mixture of left and right hemispheres across subregions within each mouse, and across mice for each subregion. During each session for each mouse, we recorded simultaneously from all 3 subregions.

**Figure 2.**
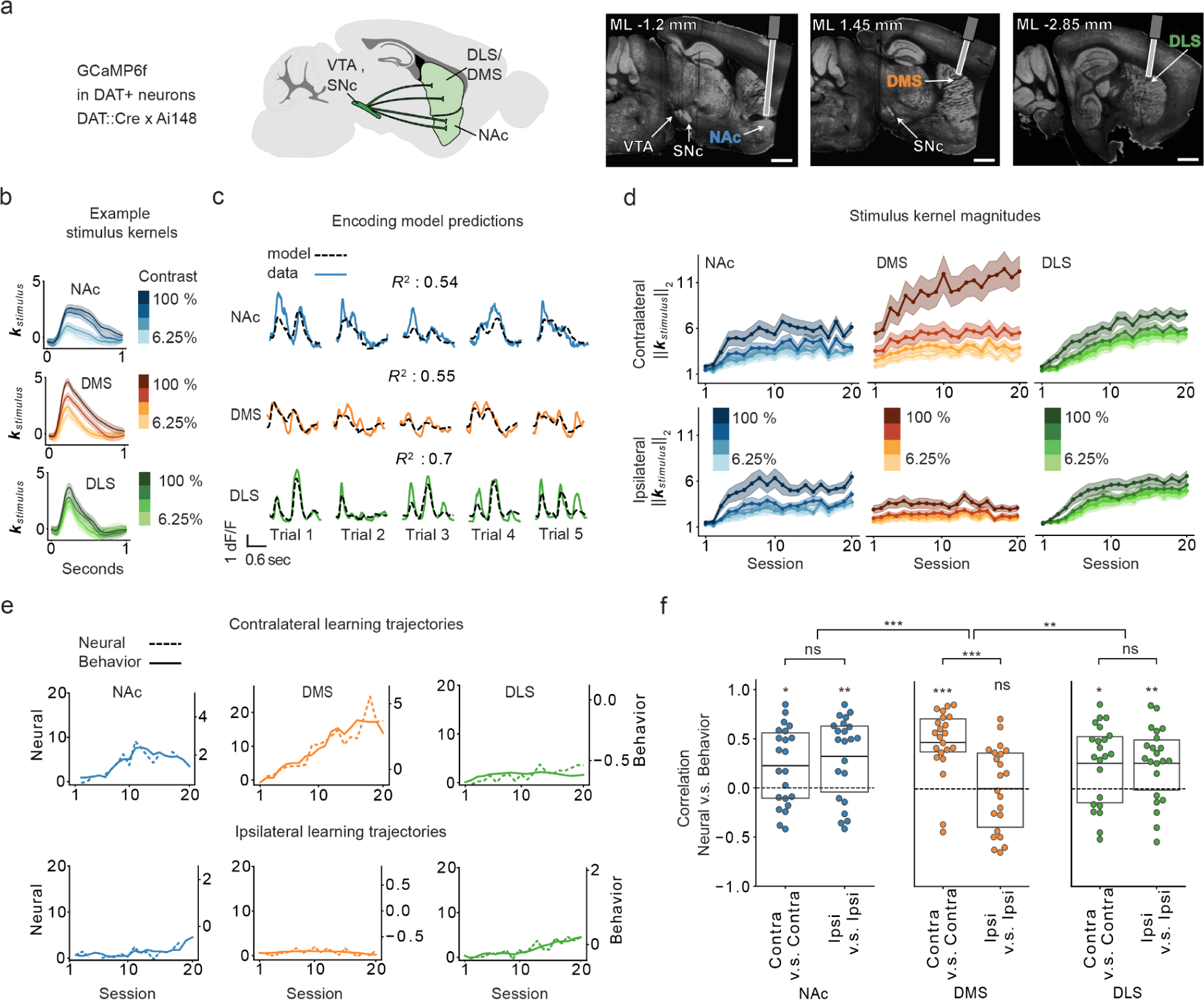
In DA terminals across striatum, contrast-dependent visual responses track individual side-specific learning trajectories. **a.** Experimental strategy used for collecting the fiber photometry data from DA terminals. Left: Schematic of the recorded projections using the GCamPG6f x DAT::Cre mouse line. Right: example histology. Scale bar: 1 mm. **b.** Contralateral stimulus response kernels, from an example mouse on an example session. **c.** z-scored dF/F (solid line) and predictions from the encoding model (dashed line) on 5 different trials for an example mouse, on an example session. *R*^2^ is the variance explained across the session within all trial epochs (from stimulus onset to 1 second after feedback). **d.** Stimulus response magnitudes (L2-norm) in each region and session, averaged across mice, for contralateral (top) and ipsilateral (bottom) stimuli. Lines and shading represent mean +/- s.e.m. **e.** Trajectories of the contrast-dependence of neural stimulus response magnitudes (“neural”; difference in L2-norm for 100% and 6.25% contrast) and the behavioral stimulus choice weights (“behavioral”), for contralateral (top) and ipsilateral (bottom) stimuli. **f.** Correlations of the neural and behavioral trajectories as shown in (e). values calculated with t-tests. * p < 0.05, ** p < 0.01, *** p < 0.001, ns: not significant. See Table 2 for statistical details for panel (f). *n*=*22* mice in (d) and (f).

**Table 2.** Statistics for Fig 2.f.

|  | p-value | t-value | df | se | n |
| --- | --- | --- | --- | --- | --- |
| **Contra correlations: test against 0** |  |  |  |  |  |
| **NAc** | 0.0140 | 2.67 | 21 | 0.084 | 22 |
| **DMS** | <1e-05 | 6.5 | 21 | 0.073 | 22 |
| **DLS** | 0.0063 | 3.05 | 21 | 0.08 | 22 |
| **Ipsi correlations: test against 0** |  |  |  |  |  |
| **NAc** | 0.0006 | 4.06 | 21 | 0.08 | 22 |
| **DMS** | 0.897 | 0.13 | 21 | 0.09 | 22 |
| **DLS** | 0.001 | 3.79 | 21 | 0.07 | 22 |
| **Contra & Ipsi comparison: test difference in means** |  |  |  |  |  |
| **NAc** | 0.357 | -0.93 | 42 | 0.11 | [22, 22] |
| **DMS** | 0.0003 | 3.92 | 42 | 0.11 | [22, 22] |
| **DLS** | 0.869 | -0.16 | 42 | 0.11 | [22, 22] |
| **Contra minus Ipsi: test difference in means** |  |  |  |  |  |
| **NAc & DMS** | 0.0004 | -3.84 | 42 | 0.14 | [22, 22] |
| **DMS & DLS** | 0.003 | -3.15 | 42 | 0.15 | [22, 22] |

Given that DA neurons encode reward predictive cues^15^, behavioral and dopaminergic sensitivity to the stimuli should be expected to increase during task acquisition, as animals learn that the stimuli are reward predictive^8,9,23–26^. To test this, we quantified dopaminergic sensitivity to the visual stimuli by fitting the normalized fluorescence data with a linear encoding model for each session and subregion (model schematic in Supp. Fig. 3). This allowed us to estimate stimulus response kernels (Fig. 2b), which reflect the contribution of the visual stimulus to the neural signals while (linearly) accounting for other task events (actions, outcomes). This model could accurately capture moment by moment fluctuations in fluorescence on each trial (Fig. 2c; Supp. Fig. 3).

Averaged across mice, across all regions, the magnitude of these stimulus responses increased across sessions, as mice learned that the stimuli were reward predictive (Fig. 2d; L2-norm of stimulus response kernels). Moreover, stronger stimulus responses emerged to the stronger contrasts, consistent with stronger contrasts becoming more reward-predictive as the animals acquired the task (increasing stimulus sensitivity behavioral weight with training in Fig. 1f-j). While stimulus responses in DLS and NAc DA were similar for contralateral and ipsilateral stimuli, DMS had much stronger stimulus responses for contralateral than ipsilateral stimuli (Fig. 2d)^28,31^.

How do these stimulus responses relate to each mouse’s idiosyncratic and side-specific learning profiles (Fig. 1)? For each mouse, across sessions, we correlated the contrast-dependence of the *behavioral* trajectories for stimuli on one side (stimulus contrast weight from the behavioral model; Fig. 1) with the contrast-dependence of the *neural* trajectories for stimuli on the same side (difference in L2-norm for highest vs lowest contrast stimulus kernel). There was a strong correlation between the behavioral and neural measures across all regions (example trajectories: Fig. 2e; neural and behavioral correlations for all mice: Fig. 2f). While DLS and NAc DA showed this correlation for both ipsilateral and contralateral stimuli, in the DMS, this correlation was only significant for contralateral stimuli (Fig. 2f), presumably reflecting the contralateral specificity of the stimulus responses themselves (Fig. 2d).

Thus, across striatum, we observed the co-evolution during task acquisition of side-specific dopaminergic and behavioral sensitivity to the visual stimuli, consistent with reward prediction error signaling in DA neurons.

### Pre-existing visual responses in DMS DA terminals predict learning on the contralateral side

While the striatum-wide correlations between DA trajectories and learning trajectories were striking (Fig. 2), they do not clarify whether there are DA signals that *precede* behavioral changes that could potentially explain individual differences in task acquisition. Notably, the DMS (unlike the other regions) exhibited contrast-dependent responses to the visual stimuli from the very first session (Fig. 2d). To determine if these signals existed before training, or if they emerged quickly on the first session, we examined DA responses during an earlier pre-exposure session (“session 0,” before the first session) in which the visual gratings were presented, but mice did not receive rewards, nor could they turn the wheel (Fig. 3a).

**Figure 3.**
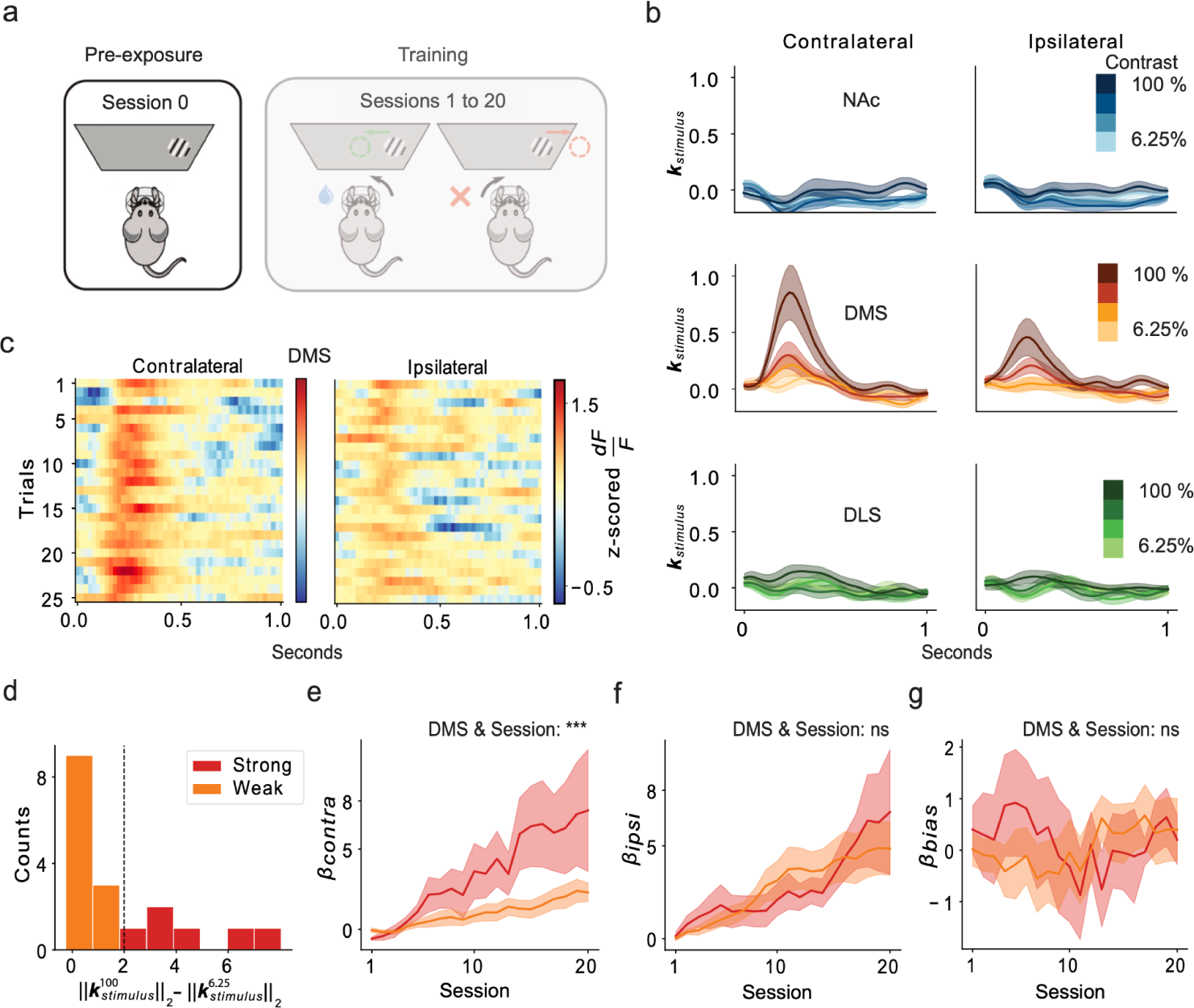
Pre-existing visual responses in DMS DA terminals predict side-specific learning trajectories. **a.** Schematic of the stimulus pre-exposure session before training (“Session 0”). **b.** Stimulus response kernels for contralateral and ipsilateral stimuli of each contrast, averaged across mice, during Session 0. **c.** Heatmap of stimulus responses on Session 0 to 100% contrast stimuli for the first 25 trials, averaged across mice. **d.** Histogram across mice of contrast-dependent stimulus responses on session 0, quantified as the difference in the L2-norm of the highest and lowest contrast contralateral stimulus, colored by weak (orange: < 2) and strong (red: > 2). **e.** Contralateral stimulus sensitivity weights from the behavioral model, for mice with strong versus weak contrast-dependent stimulus responses during session 0 (subdivision of mice shown in (d)). *** p < 0.001 for the interaction between DMS stimulus response on session 0 & session in a 2-way ANOVA (see Table 3.1-3.2 for model details and full results). **f.** Same as (e), except for the ipsilateral stimulus weight from the behavioral model. No significant interaction (ns) between DMS stimulus response on session 0 and day (see Table 3.3-3.4 for model details and full results). **g.** Same as (e,f), but for the bias weights from the behavioral model (transformed such that positive means contralateral bias). No significant interaction (ns) between DMS stimulus response on session 0 & session (see Table 3.5-3.6 for model details and full results). In all panels, *n*=18 mice.

**Table 3.1.** Statistics for Figure 3.e (individual coefficients)

$$\beta_{contra} \sim 1 + dms * session + (1 + session | mouse)$$
| Variable | Coef. | Std. Error | z | Pr(>\|z\|) |
| --- | --- | --- | --- | --- |
| Intercept | 0.348517 | 0.176679 | 1.97 | 0.0485 |
| session: middle | 1.53946 | 0.395762 | 3.89 | 0.0001 |
| session: late | 3.14228 | 0.655547 | 4.79 | <1e-05 |
| dms | 0.11926 | 0.073055 | 1.63 | 0.1026 |
| session: middle & dms | 0.44442 | 0.163799 | 2.71 | 0.0067 |
| session: late & dms | 1.22895 | 0.271316 | 4.53 | <1e-05 |

**Variable coding:** See variable coding in statistics section

**Table 3.2.** Statistics for Figure 3.e (ANOVA, type 3)

$$\beta_{contra} \sim 1 + dms * session + (1 + session | mouse)$$
| Variable | DOF | Res.DOF | F value | Pr(>\|F\|) |
| --- | --- | --- | --- | --- |
| Intercept | 1 | 337 | 3.8259 | 0.0513 |
| session | 2 | 15 | 11.5722 | 0.0009 |
| dms | 1 | 15 | 2.6203 | 0.1263 |
| session & dms | 2 | 337 | 10.2875 | <1e-04 |

**Variable coding:** See variable coding in statistics section

**Table 3.3.** Statistics for Figure 3.f.

$$\beta_{ipsi} \sim 1 + dms * session + (1 + session | mouse)$$
| Variable | Coef. | Std. Error | z | Pr(>\|z\|) |
| --- | --- | --- | --- | --- |
| Intercept | 0.912165 | 0.261839 | 3.48 | 0.0005 |
| session: middle | 1.97641 | 0.539155 | 3.67 | 0.0002 |
| session: late | 3.90767 | 0.972914 | 4.02 | <1e-04 |
| dms | 0.083602 | 0.108319 | 0.77 | 0.4402 |
| session: middle & dms | -0.27578 | 0.223162 | -1.24 | 0.2165 |
| session: late & dms | -0.181152 | 0.402711 | -0.45 | 0.6528 |

**Variable coding:** See variable coding in statistics section

**Table 3.4.** Statistics (ANOVA, type 3) for Figure 3.f.

$$\beta_{ipsi} \sim 1 + dms * session + (1 + session | mouse)$$
| Variable | DOF | Res.DOF | F value | Pr(>\|F\|) |
| --- | --- | --- | --- | --- |
| Intercept | 1 | 337 | 11.9327 | 0.0006 |
| session | 2 | 15 | 9.7095 | 0.002 |
| dms | 1 | 15 | 0.5857 | 0.456 |
| session & dms | 2 | 337 | 0.7703 | 0.4637 |

**Variable coding:** See variable coding in statistical analysis section of Methods.

**Table 3.5.** Statistics for Figure 3.g (individual coefficients)

$$\beta_{bias} \sim 1 + dms * session + (1 + session | mouse)$$
| Variable | Coef. | Std. Error | z | Pr(>\|z\|) |
| --- | --- | --- | --- | --- |
| Intercept | 0.023347 | 0.375064 | 0.06 | 0.9504 |
| session: middle | -0.126425 | 0.253667 | -0.5 | 0.6182 |
| session: late | 0.323096 | 0.423532 | 0.76 | 0.4455 |
| dms | 0.163736 | 0.155245 | 1.05 | 0.2916 |
| session: middle & dms | -0.185426 | 0.104978 | -1.77 | 0.0773 |
| session: late & dms | -0.184996 | 0.175278 | -1.06 | 0.2912 |

**Variable coding:** See variable coding in statistical analysis section of Methods.

**Table 3.6.** Statistics for Figure 3.g (ANOVA, type 3)

$$\beta_{bias} \sim 1 + dms * session + (1 + session | mouse)$$
| Variable | DOF | Res.DOF | F value | Pr(>\|F\|) |
| --- | --- | --- | --- | --- |
| Intercept | 1 | 337 | 0.0038 | 0.9508 |
| session | 2 | 15 | 1.3924 | 0.2788 |
| dms | 1 | 15 | 1.0937 | 0.3122 |
| session & dms | 2 | 337 | 1.579 | 0.2077 |

**Variable coding:** See variable coding in statistical analysis section of Methods.

In DMS, but not NAc or DLS, DA terminals had contrast-dependent visual responses during stimulus pre-exposure before training (“session 0”; Fig. 3b). These DMS DA responses were contralateral-side-specific, similar to the pattern observed throughout training (Fig. 2d). The stronger response to higher contrast stimuli may be interpreted as a salience-related signal^23,27,33,42–49^. While DA has been associated with novelty coding^23,48,50,51^, these pre-existing visual responses did not attenuate during the 25 presentations of each stimulus (Fig. 3c).

Previous theoretical accounts suggest novelty- or salience-related DA signals could provide animals with a “bonus” (or head start) in forming stimulus-reward associations^52^. Therefore, we wondered if variability in these DMS-specific pre-existing DA responses to the visual stimuli might predict differences across individuals in learning trajectories.

Indeed, we observed a striking relationship between these pre-exposure stimulus responses in DMS DA and individual differences in side-specific learning. For visualization purposes, we separated mice into 2 groups based on their pre-exposure visual responses (Fig. 3d; “strong” vs “weak” contrast-dependent contralateral DMS DA responses on “Session 0”), and plotted the trajectory of the behavioral model weights during subsequent task training in each group (Fig. 3 e-g). Over the course of training, the animals with larger pre-exposure visual stimulus responses developed larger contralateral behavioral stimulus sensitivity weights (Fig. 3e). On the other hand, the pre-exposure DMS DA visual responses were not predictive of the ipsilateral weights or the bias (Fig. 3f,g).

Thus, pre-existing visual responses in DMS DA predict individual differences in side-specific learning, even before any training. While NAc and DLS did not have visual responses during pre-exposure (Fig. 3b), we wondered how early in training their visual responses predicted final performance, and how this compared to DMS. For each region on each training session, we computed the across-animal correlations between the DA response to visual stimuli on each side with the behavioral stimulus weights late in training (sessions 16-20) on the same side (Supp Fig. 4). These correlations appeared much earlier in training in the DMS than the other regions (Supp Fig. 4; for contralateral stimuli only), in line with pre-existing contralateral visual responses being a unique property of this population.

**Figure 4.**
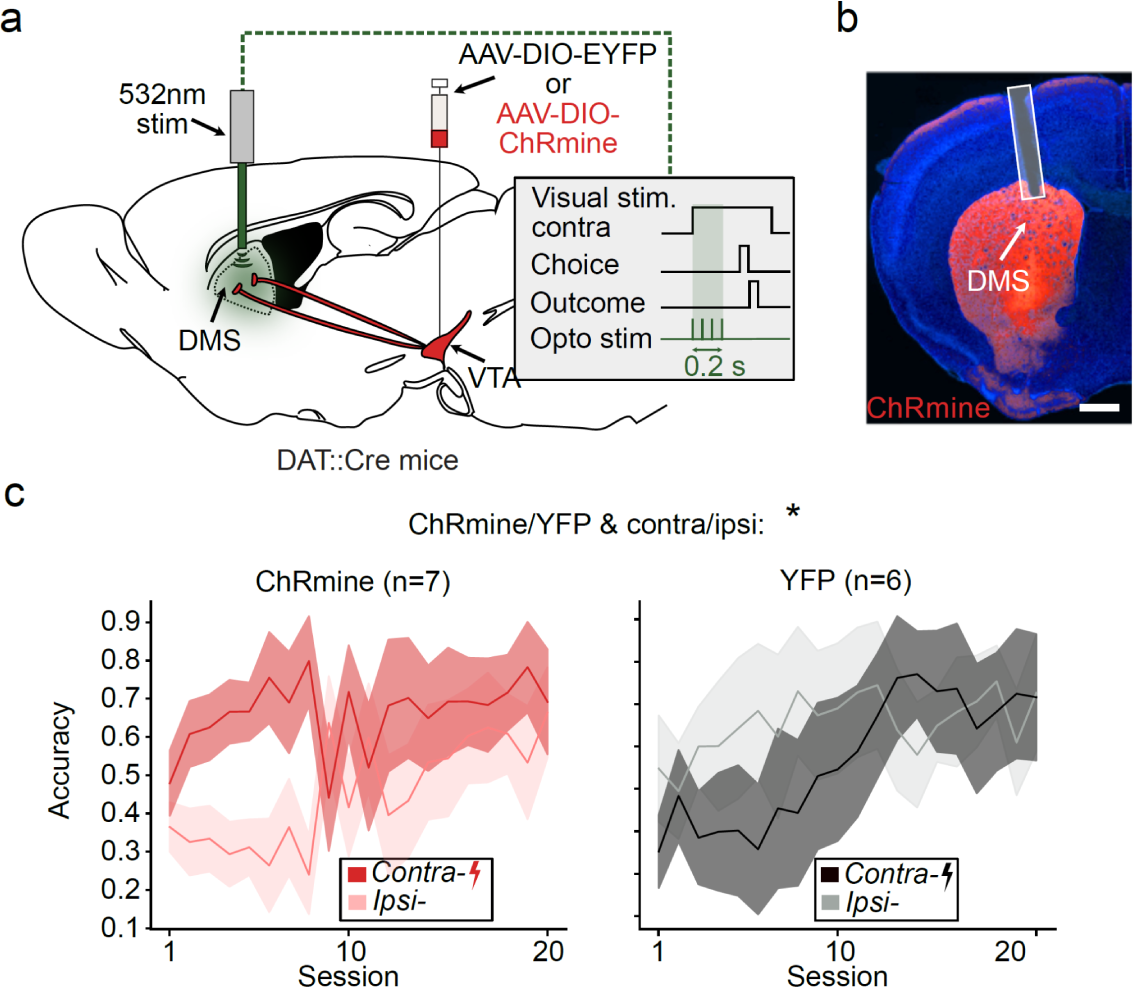
Stimulating DMS DA terminals at the onset of contralateral stimulus presentation improves side-specific performance. **a.** Schematic of the optogenetic stimulation of DMS dopamine terminals. Mice either expressed Chrmine or a control construct in DA neurons. DA terminals in the DMS were optogenetically stimulated unilaterally (532 nm, 0.2s burst duration, 5ms pulse width, 20 Hz pulses, ∼0.25 mW) at the onset of the contralateral stimulus presentation throughout training. **b.** Example histology image of optical fiber location and terminal expression of ChRmine-mScarlet. Scale bar: 900 µm. **c.** Comparison of performance for contralateral and ipsilateral stimulus trials in control (*n*=7, left panel) and ChRmine (*n*=6, right panel) mice. *p<0.05 for cohort (ChRmine/YFP) & side (contra/ipsi) interaction in 3-way ANOVA with cohort (ChRmine / YFP), day and side (contralateral / ipsilateral) as factors (see Table 4.1-4.2 for model details and full results).

**Table 4.1.**
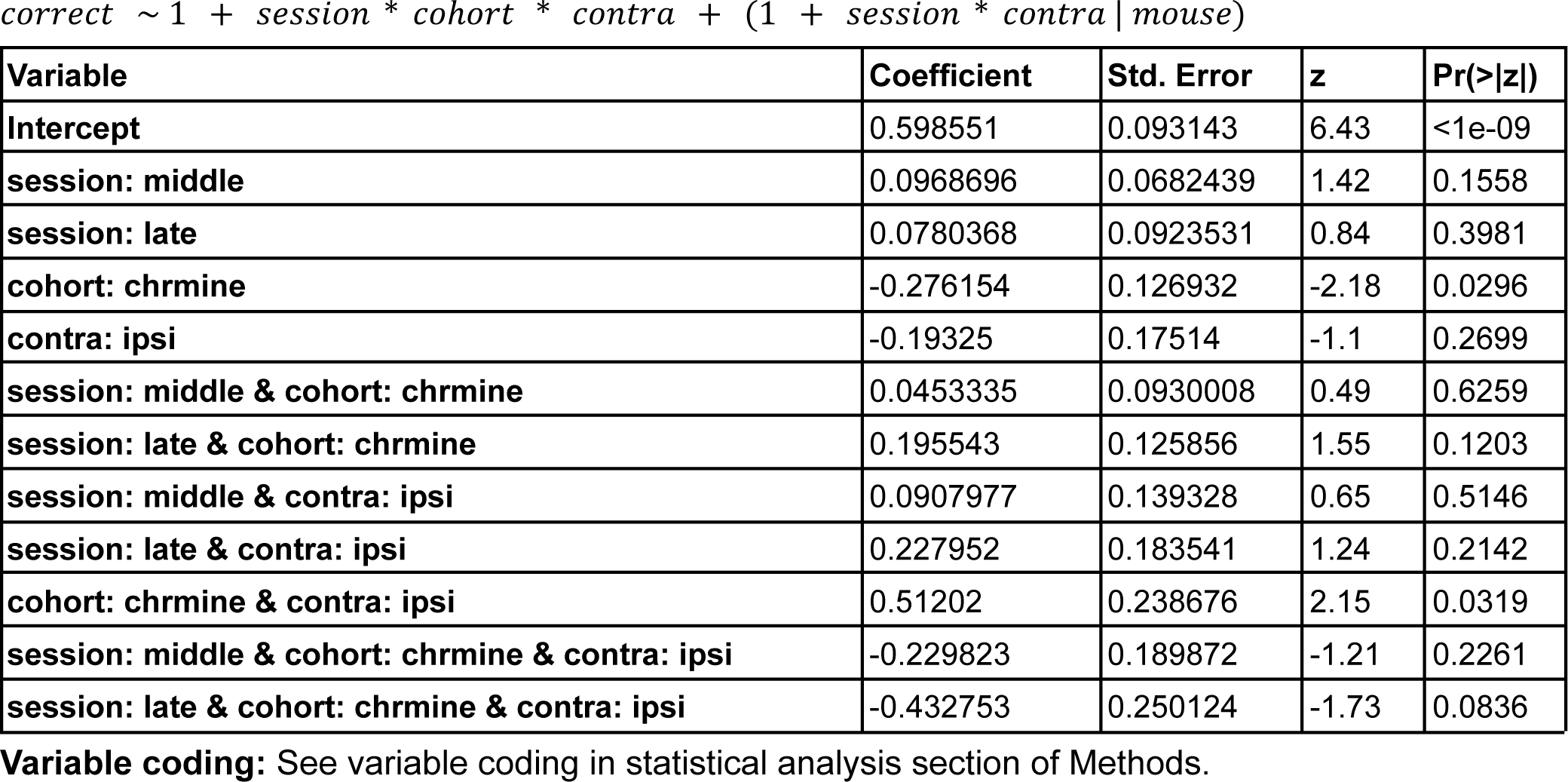
Statistics for Figure 4.c (individual coefficients)

**Table 4.2.** Statistics for Figure 4.C (ANOVA, type 3)

*correct ~ 1 + session \* cohort \* contra + (1 + session \* contra | mouse)*
| Variable | DOF | Res.DOF | F value | Pr(>\|F\|) |
| --- | --- | --- | --- | --- |
| Intercept | 1 | 498 | 40.3425 | <1e-09 |
| session | 2 | 10 | 0.9958 | 0.4033 |
| cohort | 1 | 10 | 4.624 | 0.057 |
| contra | 1 | 498 | 1.1894 | 0.276 |
| session & cohort | 2 | 498 | 1.2345 | 0.2919 |
| session & contra | 2 | 498 | 0.7554 | 0.4703 |
| cohort & contra | 1 | 498 | 4.4959 | 0.0345 |
| session & cohort & contra | 2 | 498 | 1.5 | 0.2241 |

**Variable coding:** See variable coding in statistical analysis section of Methods.

### Activation of DMS dopamine terminals during contralateral stimulus presentation improves side-specific performance

The strength of pre-existing visual responses in DMS dopamine terminals predicted side-specific learning (Fig. 3d-e, Supp Fig. 4). Could these visual signals causally impact side-specific performance during learning? While it is clear that DA at the time of outcome reinforces previous actions^53–57^, whether DA signals at the time of a preceding cue impact learning is less clear^49,53,54,58–60^.

To test for a causal role of visual-stimulus-related DA signals in DMS, we performed brief unilateral optogenetic stimulation of DMS dopamine terminals at the presentation of the contralateral visual stimulus throughout training (Fig. 4a-b, Supp Fig. 2d). The stimulation, which terminated before the outcome period (Supp. Fig. 1d-e), led to a significant improvement in performance between contralateral versus ipsilateral trials for the opsin vs control groups that grew over early training (Fig. 4c). We therefore concluded that DA signals in DMS could improve performance for contralateral stimuli.

## Discussion

As mice learn to perform a visual decision-making task, variation across individuals in learning trajectories closely tracks visual responses in DA terminals across striatum (Fig 1,2). In contrast to these striatum-wide patterns, prior to any rewards or training, pre-existing side-specific visual responses are present specifically in DMS DA terminals, and these signals predict and help explain side-specific learning trajectories (Fig 3,4). This work is significant in suggesting that i) the initial conditions of the DA system are important in explaining individual differences in learning, and ii) feature- and projection-specific DA signals could be a mechanism to simplify the problem of initial task acquisition.

### Pre-existing visual responses in DMS-projecting dopamine neurons could serve to simplify initial task acquisition

A major reason initial task acquisition is challenging is the issue of credit assignment: in virtually any task, even nominally simple ones, there are multiple possible dimensions of the environment that an animal could try to learn about, with most being reward-irrelevant. In the case of the visual decision-making task used here, mice need to learn that the side-specific relationship between visual stimuli and actions is what leads to reward, while other stimuli, actions or stimulus-action relationships do not (including many high dimensional, uncontrolled incidental features of the environment^37^).

Our data suggest that pre-existing, side-specific and projection-specific visual DA responses could serve to decrease the dimensionality of this learning problem. In particular, different striatal subcircuits could be biased to more quickly learn reward associations for different subsets of stimuli or actions: in the case of DMS, our data reveals a pair of such circuits, selective for visual stimuli that are salient and presented on one side of the visual world but not the other. Extended to other subcircuits and task features, such an architecture would effectively lower the dimensionality of the initial learning problem, as each subcircuit could be biased towards learning about a different subset of features. Indeed, other striatal circuits likely have pre-existing DA sensory responses to other modalities that could in turn predispose learning in favor of those associations. For example, pre-existing auditory responses in DA neurons in the tail of the striatum^61^ could support auditory-motor learning^62,63^, much like the visual responses we examine here in DMS.

This framework helps in interpreting the function of salience-related signals that have been reported in previous DA recordings^23,27,33,42–49^, and is consistent with a classic idea about the role of salience signals in providing an optimistic “bonus” to support learning^52^. However, it adds to this framework by suggesting there are a number of separate circuits rather than a single, global prediction error.

Indeed, such hypothetical specialization simplifies the curse of dimensionality insofar as any given subcircuit would be initially biased to quickly discovering simple associations, appropriate for tasks in which a small set of stimuli are associated with reward. Of course, there is no free lunch. Such a bias for sparse solutions would not help, and might indeed hinder, detecting contingencies that depend on more complex (e.g., multimodal) combinations of features. This reasoning also leads to testable predictions about which types of contingencies are most easily learnable: those that well-match the feature-selectivity of a projection-defined DA population. This may explain why, in the present data, animals often learn more quickly to respond to stimuli on one side or the other, independently from those on the other side, rather than the fuller response rule combining both sides (Fig. 1).

### Initial conditions of the DMS DA system can explain individual differences in task acquisition

Our data also help to explain why different individuals learn the same task much more quickly than others (Fig. 1). The variation across individuals in the pre-existing visual response in DMS DA signals, before task training and the presentation of any rewards, predicts individual differences in acquisition of the visual-motor task on the contralateral side (Fig. 3). This supports the idea that the pre-existing DMS DA visual responses facilitate reward learning, and highlights the importance of the initial conditions of the DA system in understanding the emergence of individual behavioral differences.

While DMS appears unique in having pre-existing visual responses, all subregions we examined have DA visual responses that emerge with learning and closely track learning trajectories (Fig. 2). This is consistent with the widespread encoding of a reward prediction error in dopamine terminals across the striatum^64^ because in this task the visual stimuli predict reward, and higher contrast indicates that reward is more likely (with reward expectation depending on the individual time-varying psychometrics curves, i.e. how the animal leverages the stimuli to drive correct behavior; Fig. 1).

### Relationship to prior experimental work on the DMS DA system

Our results complement prior work characterizing the DMS DA system, demonstrating contralateral response preferences^28,31,65^, visual responses^31,61^, and stimulus-value related responses^64^. Our results also relate to recent studies on striatal DA in initial acquisition of sensory decision-making tasks^8,9^, most directly the demonstration that DA responses and behavioral sensitivity to task stimuli grow together during learning^9^ (similar to our Fig. 2). Our work adds to the literature primarily by revealing a pre-existing sensory feature-specific and projection-specific DA signal that explains later learning (Fig. 3,4).

On the other hand, the relationship between our work and recent work implicating DMS DA in individual differences in the development of a habitual^17^ or punishment-resistant^21^ behavioral strategy is not yet clear. While it seems possible that a pre-existing sensory feature-specific DA signal could contribute to the emergence of those strategies, more work is required to clarify a potential connection.

While here we focus on visual responses in DMS DA, previous work from our group and others has identified putative action correlates in this population^28,65–67^. Lateralized action-related DMS DA responses may relate to orienting behavior, as DMS is a target of frontal orienting fields^68,69^, has been implicated in tasks with orienting behavioral outputs^70–72^, and the DMS DA signal itself reserves with changes in orientation^65^. For this reason, in the present task, action responses may be at least partially obscured by head-fixation which prevents orienting behavior. Regardless, we are confident the contrast-dependent responses examined here are visual, as we isolated them from wheel movement with an encoding model, and furthermore confirmed their presence during stimulus pre-exposure in the absence of task-related movements (Fig. 3a,b).

### Relationship to the feature-specific RPE model of DA heterogeneity

The presence of a visual feature-specific DMS DA signal is consistent with a recent model that proposes that response variation across DA neurons can be explained at least in part by differences in the feature representations in the inputs that are used to calculate reward prediction errors (“feature-specific reward prediction error model”^37^). In this framework, different dopamine neurons calculate different reward prediction errors based on different subsets of the full feature space, based on the corticostriatal inputs they preferentially receive. This model thus predicts similar feature selectivity of the DMS-projecting dopamine neurons relative to the DMS neurons themselves, assuming an anatomical arrangement where DMS projects (indirectly or directly) primarily to DMS-projecting DA neurons^73^. Consistent with this prediction, DMS receives direct visual cortical inputs^74^, and has visual responses before task training^75^.

### Summary

In summary, we find that the presence of pre-existing visual responses in DMS DA provides a simple explanation for why some individuals learn a visual decision-making task faster than others. This discovery sheds light on the significance of DA neuron functional heterogeneity: it could serve to bias different target regions towards initially learning about different subsets of task features, which could help address the dimensionality of the initial task learning problem. This concept generates concrete, testable predictions about which reward associations are easily learnable: those with stimuli that match the pre-existing responses of a projection-defined DA population.

## Methods

### Animals

For the fiber photometry experiments (Fig. 1-3), a total of 22 mice (*n=14* male and *n=8* female) were used from a cross of DAT::IRES-Cre mice (JAX 006660) and the GCaMP6f reporter line Ai148 (JAX 030328). For the optogenetic experiments (Fig. 4), we used a total of 13 DAT::IRES-Cre mice (*n=4* male and *n=9* female). Mice were maintained on a reversed 12 h light cycle and experiments were formed on their dark cycle. All mice used were 3-4 months old at the start of training. All experimental procedures were conducted in accordance with guidelines from the National Institutes of Health and were reviewed by the Institutional and Animal Care Use Committee at Princeton University.

### Surgery

Prior to the start of the surgery, mice received a preoperative antibiotic (5 mg/kg Baytril) and analgesic (10 mg/kg Ketofen). Postoperative analgesic (10 mg/kg Ketofen) was administered daily for 3 days from the day of the surgery.

#### Headbar implantation

For all stereotaxic surgeries, mice underwent sterile stereotaxic surgery under anesthesia (5% isoflurane for induction, 1.5-2% for maintenance). Briefly, the scalp and underlying periosteum was removed. Bregma and lambda were leveled, and a small steel headbar was centered at -6.9 mm Anterior-Posterior relative to bregma and cemented to the skull with Metabond (Parkell). Headbar implantation was followed by virus infusion and/or optical fiber implantation (see sections below).

#### Optical fiber implantation

For fiber photometry experiments (data shown in Fig. 1-3), low-autofluorescence optical fibers encased in a ferrule (0.37 NA, ⌀200 µm core, 1.25mm ferrule, Neurophotometrics) were implanted at each of the following locations (fiber tip location relative to bregma):

- DLS: 2.6 mm (Medio-lateral, M-L), 0 mm (Anterior-Posterior, A-P), -2.8 mm (Dorso-ventral, D-V).
- DMS: 1.5 mm M-L, 0.74 mm A-P, -2.4 mm D-V.
- NAc: 1 mm M-L, 1.45 mm A-P, -4.5 mm D-V.

In each mouse, fibers targeting DLS and NAc were always inserted in the same hemisphere, the DMS fiber was positioned in the opposite hemisphere. The hemisphere allocation was counterbalanced across mice. For DLS, the location above was targeted with a fiber rotated at 10° in the M-L/D-V plane. For the optogenetics experiments (Fig. 4), fiber optic fibers (⌀300 µm core/0.39 NA, 2.5 mm ferrule, ThorLabs) were implanted bilaterally to target the DMS at the following coordinates: +/- 1.5 mm M-L, 0.74 mm A-P, -2.4 mm D-V. These locations were reached with a 10° M-L/D-V rotation..

#### Virus injections

For the optogenetics experiments (Fig. 4), AAV2/5-EF1a-DIO-ChRmine-mScarlet-WPRE-hGHpA (opsin virus, titer: 9e^12^ genome copies/ml, Princeton Neuroscience Institute viral core) or AAV2/5-EF1a-DIO-EYFP-WPRE-hGHpA (control virus, titer: 1.5e^14^ genome copies/ml, Princeton Neuroscience Institute viral core) was infused bilaterally in the VTA-SNc (+/- 1 mm M-L, -3.1 A-P, -4.66 D-V) of ∼4-6 weeks old mice. 500 nl were infused in each hemisphere at a speed of 75 nl/min. In order to achieve sufficient terminal expression by the start of training (3/4 months), all viral injections were performed a minimum of 8 weeks in advance of training and prior to the headplate implantation surgery.

### Behavioral task

#### Behavioral apparatus

We used the standardized behavioral apparatus from the International Brain Laboratory. For detailed instructions on the components and operations of behavioral apparatus used please see ^76^. Briefly, the rig consisted of an LCD screen (LP097Q × 1, LG) and a custom 3D-printed mouse holder and head fixation system that held the mouse in front of the screen such that its forepaws rested on a rubber steering wheel (86652 and 32019, LEGO). A spout was positioned in front of the holder, which the mouse could reach it with its tongue but it did not occlude the field of vision. The spout was connected to a water reservoir and water flow was controlled with a solenoid pinch valve (225P011-21, NResearch). The rig was constructed with Thorlabs parts inside a small soundproof cabinet (9U acoustic wall cabinet 600 × 600, Orion). A speaker (HPD-40N16PET00-32, Peerless by Tymphany) positioned on top of the screen was used to play task-related sounds, and an ultrasonic microphone (Ultramic UM200K, Dodotronic) was used to record ambient noise from the rig. Wheel position was recorded with a rotary encoder (05.2400.1122.1024, Kubler) controlled by the Bpod Rotary Encoder Module (Sanworks). Video of the mouse was recorded with a USB camera (CM3-U3-13Y3M-CS, Point Grey). All task-related devices were controlled by a Bpod State Machine (Sanworks) and synched with a data acquisition board (USB201, Measurement Computing). The task logic was programmed in Python and the visual stimulus presentation and video capture was handled by Bonsai^77^ and the Bonsai package BonVision^78^.

#### Behavioral task and training

We used a standardized visual decision-making task^7,38^. In this task, mice are head fixed in front of a LCD screen. A visual grating (Gabor patches, 0.1 spatial frequency) whose contrast varied across trials (100%, 50%, 25%, 12.5%, 6.25%) appeared on either the right or left side of the screen (+/- 35° azimuth), accompanied by a 0.1 s tone (5 kHz sine wave, 10ms ramp). A steering wheel that could be used to move the visual grating along the horizontal axis was placed under the mouse’s paws (4° of visual grating movement /mm of wheel movement). The mouse could obtain a small reward of 10% sucrose water (3 μl) by moving the visual grating to the center of the screen. On the contrary, if the mouse steered the grating away from the center (35° from initial position) or failed to center the grating in 60 s, the trial was considered an error. Errors were signaled by the lack of reward delivery and a brief noise (0.5 s, 65 dB, white noise). After a choice was completed (correct or incorrect), wheel movements could no longer move the visual grating for 1 or 2 seconds on correct versus incorrect trials, respectively. After this timeout, all trials were followed by a 0.5 s inter trial interval where no gratings were presented. In order for a new trial to start, the steering wheel had to be still for a “quiescent period”, whose duration was randomly sampled on a trial by trial basis from an exponential distribution of mean 0.55 s and truncated from 0.4 to 0.7 s.

To simplify the interpretation of neural activity correlates of behavior across learning, mice experienced the full extent of the task (no shaping or debiasing protocol) from the first session. All mice were trained for a minimum of 18 sessions, 5-7 days a week, for a 1h session each day. In order to motivate the mice to do the task, mice had restricted water access from 1 week before starting training until the end of the experiment. We monitored that their weight never dropped more than 20% from their pre-water restriction weight, and ensured that they consumed a daily minimum of 1 ml of water per 25 g of weight. Most mice were able to obtain their daily allocation of water through the task alone after a few sessions. When this minimum was not achieved, mice were supplemented at the end of the day. Mice did not have a limit on how much water they could obtain in the task (See Supp. Fig. 1a for average trials completed across training). Mice were video recorded every session.

#### Stimulus pre-exposure

For the experiment in Fig. 3, before training on the task (1-10 days prior to the start of training), 18 mice underwent two 1h pre-exposure sessions where we measured neural responses to task features in the absence of reward. In the first pre-exposure session, mice were presented with the same visual gratings used in the task (with the same range of contrasts) on either side of the screen for 250-272 trials with a 10 seconds inter-trial interval. The stimulus contrast and side in each trial was randomly sampled between the 8 possible combinations. As in the standard task, presentation of the visual gratings was accompanied by a brief 0.1 s tone. However, during this pre-training session the wheel was locked and the visual gratings remained static on either side of the screen. In the second session (data not shown), mice were allowed to move the wheel and move the visual grating but no rewards were given.

### Fiber photometry

#### Data acquisition

We simultaneously recorded GCaMP6f signals from DA terminals in DMS, DLS and NAc with a multi-fiber photometry system (FP3002, Neurophotometrics) controlled with the Bonsai Neurophotometrics module^79^. Briefly, the system consists of a CMOS camera acquiring fluorescence emissions and an LED exciting with 470 nm light of 10 ms width pulses at 50Hz (464/476 sessions) or 20 Hz (12/476 sessions). At the tip of the patch cable, the excitation light was ∼0.4 mW. The camera acquisition epochs were timed with the emission lights. We used a Low Autofluorescence Patch Cord with 3 branches (BBP(3)_200/220/900-0.37_2m_SMA-3xMF1.25_LAF, Doric) to be able to image DMS, DLS and NAc simultaneously. Prior to each recording day, we passed 0.5 mW 470 nm light through the patch cord for 1 hour in order to photobleach autofluorescence within the patch cord, and improve recording quality.

#### Signal processing

Fluorescence signals recorded during each session from each location were transformed to dF/F using the following formula:

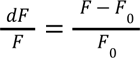

*F*_0_ was the +/- 30 s rolling average of the raw fluorescence signal. Finally, dF/F signals were z-scored per-session, using a mean and standard deviation calculated based on all the data from each session. To be included for analysis, every recording (i.e. one session from one fiber location) had to have at least one >=1% dF/F & > 3 standard deviation transient for every 10 min of recording (55/1440 recordings excluded). All data were sampled or resampled at 50 Hz for analysis.

### Histology

To confirm the locations of the opticals fibers and viral expression (Supp. Fig. 2), mice were anesthetized with pentobarbital sodium (2 mg/kg, Euthasol) and transcardially perfused first with 10 ml of ice-cold phosphate buffered saline (PBS) followed by 25 ml of 4% paraformaldehyde (PFA) in PBS. Brains were then dissected and post-fixed in 4% PFA overnight at 4°C. After fixation, brains were sliced with a vibrating blade microtome (Vibrotome VT100S, Leica) and mounted with DAPI Fluoromount-G (Southern Biotech). All slices were imaged with an automated slide scanner (NanoZoomer S60, Hamamatsu).

### Optogenetic stimulation

For the optogenetic experiment in Fig. 4, fibers were implanted bilaterally in the DMS to avoid potential behavioral biases related to an asymmetrical surgery. Stimulation was delivered unilaterally to DMS terminals expressing the red-shifted opsin ChRmine. The stimulated hemisphere was chosen randomly and kept constant throughout training for any given animal. The group identity of the mice (opsin vs control) were blinded to the experimenter throughout the duration of training. The stimulation procedure consisted of a 200 ms laser train timed to the onset of any visual stimulus presentation contralateral to the stimulated hemisphere. Each 200 ms train of stimulation consisted of 20 Hz and 5 ms width light pulses at a wavelength of 532 nm (Shanghai Laser and Optics & Co). The light power was adjusted daily to 0.25 mW at the fiber tip (in the brain). Light power was chosen to ensure activation of the terminals immediately below the fiber tip but minimize off-target activation outside DMS. ChRmine can reliably be activated with an irradiance of >= 0.1 mW/mm^2^ ^80^. Therefore, we chose a stimulation power that ensured irradiance above this threshold within but not outside of DMS. According to ^81^ the chosen power and fiber (∼0.25 mW 532nm light through ⌀300 µm core/0.39 NA fiber) yields an irradiance of 0.88 mW/mm^2^ (above threshold) just below the fiber tip (DMS) and 0.01 mW/mm^2^ (below threshold) at the DMS/NAc border (1.7 mm ventral from the fiber tip).

### Behavioral model

Our approach in modeling behavior aims to descriptively characterize the relatively long time-scale dynamics of learning that would be required to correctly associate stimuli, actions, and outcomes, particularly in the absence of shaping, de-biasing, or other experimental protocols. This relates to previous modeling efforts of similar datasets; however, instead of focusing on trial-to-trial fluctuations in psychophysical weights^82^ or the emergence of multi-state behavior^83^, we focus on session-level changes in psychophysical weights. We leveraged advances in MCMC^84–86^ to infer a set of parameters and weights for Bernoulli generalized linear models (GLM) that were expressive enough to capture the full set of behaviors that mice in our task explored.

To model the behavioral data, we built a hierarchical Bernoulli GLM to describe the relationship between the animal’s choices and a variety of task covariates. The dependent variable was per-trial choice (a Bernoulli variable). The covariates included the stimulus presented to the animal on each trial (capturing the classic psychometric curve) together with two additional effects: the animal’s exponentially filtered choice history, and a side-specific bias. We parameterized the stimulus using two regressors, x_L_ and x_R_, corresponding to the contrast of the left-side and right-side stimulus on each trial; because the stimulus only appeared on a single side in each trial, one of these regressors was zero on each trial. We transformed each contrast regressor using a tanh function: 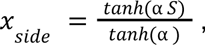 where *S* is either X_L_ or X_R_ and α is a positive constant governing the shape of the nonlinear transformation. Dividing by *tanh*(α) ensures that at 100% contrast trials *x_side_* = 1. This parametrization allowed the model to saturate at contrast levels below 100%, sidestepping the need to use lapse parameters to account for the flattening of the psychometric function at high contrast levels ^87–89^. We generated the choice history regressor *c_t_* by exponentially filtering previous choices with time constant π: *c*_*t*_ = *c*_*t*−1_ + π (*y*_*t*−1_ − *c*_*t*−1_), where choice *y_t_* takes values of -1 and +1 for left and right choices, respectively. We inferred the time constant π using MCMC sampling, along with the other model parameters. (See below.)

We took the parameters governing this model (see table below) as constant within each session and mouse. For each mouse we built a hierarchical model over sessions, instantiating a separate set of choice parameters for each session. We placed broad prior distributions on the means of the choice weights on the first session. For all subsequent sessions, we assigned the choice weights a prior centered around the previous session’s inferred values.

In particular, choice weight priors for session *d* took the form of a Student’s t-distribution: β_*d*_ ∼ *StudentT*(ν, β_*d*−1_, Σ), where ν is the degrees-of-freedom parameter for the Student T prior. This allowed the model to partially pool data across sessions (smoothing the estimates), but to do so adaptively per-animal, taking on larger values of ν for animals that steadily and slowly learned the task, and smaller values for ν for animals that had sudden and large changes in their learning. We also inferred the covariance of the StudentT priors, allowing parameters to change across sessions in coupled fashion. Unlike the choice weights, we used a single shared covariance across all sessions. This covariance was parameterized as the quadratic-form product of a diagonal matrix: *D* = σ ⊙ *I* and a correlation matrix Ω: Σ = *D*Ω*D*. The diagonal had a truncated Gaussian prior. The correlation matrix Ω was constructed from a lower triangular matrix *L*, which is a cholesky factor of the correlation matrix. These factors had a prior distribution *LKJCholesky*. The *LKJCholesky* prior itself has a parameter that tunes the strength of the correlations of the cholesky factor, which we also inferred. Functions and distributions specified here were from the STAN probabilistic programming language, and all model fits were performed in STAN^90^.

We summarize the behavioral model below, (we note below that *x*_*side*_ refers to only the contrast regressors, while *x* refers to the vector of all regressors e.g. intercept, contrast, choice history):

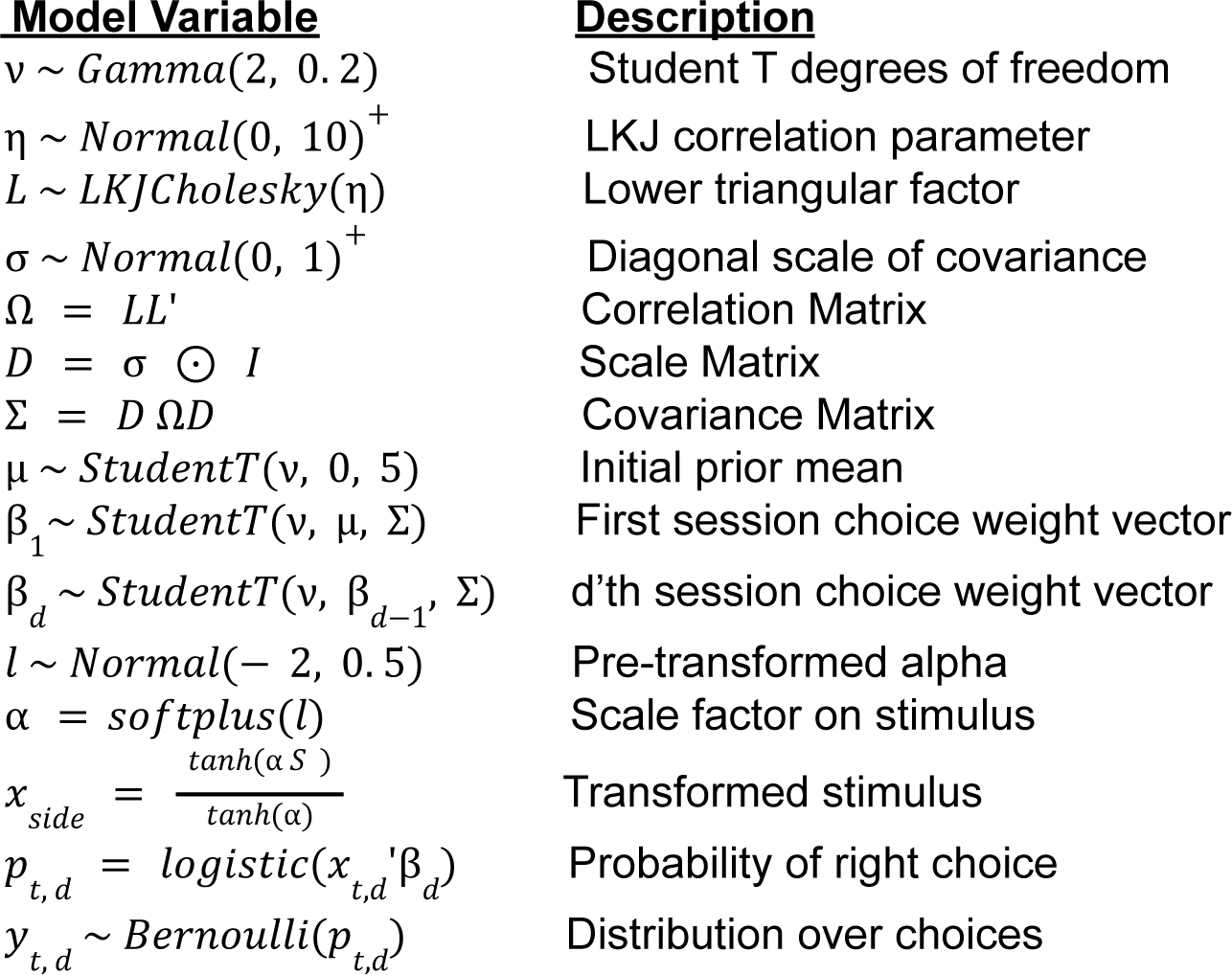

| **<u>Model Variable</u>** | **<u>Description</u>** |
| --- | --- |
| $\nu \sim \text{Gamma}(2, 0.2)$ | Student T degrees of freedom |
| $\eta \sim \text{Normal}(0, 10)^+$ | LKJ correlation parameter |
| $L \sim \text{LKJCholesky}(\eta)$ | Lower triangular factor |
| $\sigma \sim \text{Normal}(0, 1)^+$ | Diagonal scale of covariance |
| $\Omega = LL'$ | Correlation Matrix |
| $D = \sigma \odot I$ | Scale Matrix |
| $\Sigma = D\Omega D$ | Covariance Matrix |
| $\mu \sim \text{StudentT}(\nu, 0, 5)$ | Initial prior mean |
| $\beta_1 \sim \text{StudentT}(\nu, \mu, \Sigma)$ | First session choice weight vector |
| $\beta_d \sim \text{StudentT}(\nu, \beta_{d-1}, \Sigma)$ | d'th session choice weight vector |
| $l \sim \text{Normal}(-2, 0.5)$ | Pre-transformed alpha |
| $\alpha = \text{softplus}(l)$ | Scale factor on stimulus |
| $x_{side} = \frac{\tanh(\alpha S)}{\tanh(\alpha)}$ | Transformed stimulus |
| $p_{t,d} = \text{logistic}(x_{t,d}'\beta_d)$ | Probability of right choice |
| $y_{t,d} \sim \text{Bernoulli}(p_{t,d})$ | Distribution over choices |

### Neural model

To model the dopaminergic signals across learning, we built a linear-Gaussian regression encoding model to describe the relationship between task events such as the visual stimuli, actions, and reward delivery with the measured dopamine (DA). Since these events can be correlated in time and their effects on DA are partly overlapping, estimating such an encoding model helps to tease apart their individual contributions.

The regression model was defined by a set of temporal kernels that describe the DA impulse response to different task-related events, namely “stimulus”, “action”, and “reward”. For stimulus onset events, we used contrast-specific right and left temporal kernels, giving us 4 temporal kernels per side. All kernels were strictly causal, lasting for a period of 1 second (50 Hz).

Similarly, we used contrast-specific action kernels triggered at the onset of the first significant wheel movement for left and right choices (first movement larger than 0.1 radians after the end of the quiescent period). In addition to separating these kernels by contrast and side (right / left choices) we separated them by correct and incorrect trials, resulting in 8 temporal kernels per side. Separating the action kernels in this manner provided an estimate of the DA response to the interacting effects of initial stimulus location and the movement of the stimulus towards or away the center of the screen. Finally, we defined reward kernels corresponding to the moment when the animal received a water reward or a short time-out period in the same fashion as the action kernels, giving us another 8 temporal kernels per side. Thus, in total we had 40 temporal kernels in the encoding model.

We parameterized the temporal kernels in this model using a basis of linearly scaled “raised cosine” functions spanning a 1-second window after each event ^91^. The cosine basis significantly reduces the dimensionality of the design matrix *X* (compared to a full series of individual lagged event dummies). The effect on estimation of using a cosine basis is regularization. Use of a temporally smooth basis is also justified by the observation that temporally adjacent responses are strongly correlated.

We used ridge regression to estimate the model parameters, with ridge parameter γ and observation noise 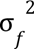 estimated via evidence optimization ^92^. We optimized for the vector of weights β, and the two scalars 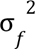, γ, which are related to the vector of neural response *f* as follows: 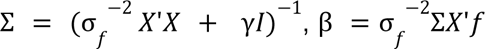

All weights that made up the entire set of temporal kernels were denoted by β, and could be indexed by their corresponding event type. For example the vector of weights β_1:50_ contained the weights for the temporal kernel for stimulus appearing on the right at 6.25% contrast (after one transforms them into the standard basis). The vector of weights β_51:100_ contained the weights for the temporal kernel for stimulus appearing on the right at 12.5% contrast, and so on for all remaining contrast levels, and event types. We further computed summary statistics of these temporal kernels, specifically the L2-norm for the stimulus responses. These summary statistics gave us a scalar measure of neural response for each training session that we then related to the estimates from the Bernoulli GLM.

The encoding model for figure 3 was fit as described above, however we only modeled the stimulus responses, and so the full set of coefficients that made up the kernels was restricted to an intercept, the 4 temporal kernels for stimulus appearing on the right, and 4 temporal kernels for stimulus appearing on the left side of the screen.

### Statistical analysis

All statistics reporting a correlation coefficient and a p-value on that correlation coefficient were computed using robust regression, in order to reduce the sensitivity of our statistical conclusions to outliers. Robust regression was performed using the *rlm* function from the RobustModels package in the Julia programming language. We used MM-estimators with a Geman Loss ^93^. For the robust regressions, we computed correlation coefficient-like statistics analogous to Pearson’s R for classic regression. In particular, we computed a pseudo- *R*^2^ statistic, and its signed square root *r*, using the RobustModels package *deviance* and *nulldeviance* functions: 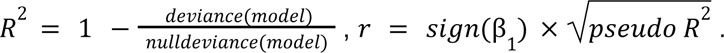 Deviance is a generalization of the residual sum of squares for linear models, and null deviance is a generalization of the total sum of squares.

Statistics in Fig. 2f were computed with the *OneSampleTTes*t, and *EqualVarianceTTest* functions in the HypothesisTests package from the Julia programming language. Significance was determined at p < 0.05, and all p-values reported are two-sided unless otherwise noted. See Table 2 for detailed results of these tests.

Statistical tests for group differences in behavioral trajectories in Fig. 3e-g and 4c, were carried out with the MixedModels and AnovaMixedModels packages in the Julia programming language. Linear Mixed Models from the MixedModels package were used to test simple effects such as the relationship between session 0 DMS stimulus response on behavioral weight values within each training period (early, middle, late). We further used type-3 F-test ANOVAs from the AnovaMixedModels package to test the overall effects in the model, such as, across training periods, is there an influence of session 0 DMS strength on behavioral weight trajectories. For all Linear-Mixed Models and ANOVAs, a*b*c expands into a + b + c + a*b + a*c + b*c + a*b*c.

#### Linear Mixed Models variable coding

Across tables 3.1 to 4.2, the variable “session” is a transformation of sessions 1 to 20. Sessions are split into 3 categories: early, middle, and late. The early category contains sessions 1 to 7, the middle category contains sessions 8 to 14, and the late category contains sessions 15 to 20. This categorical coding of sessions is motivated by the non-linear trajectory of accuracy in Fig. 4c. LinearMixedModels package in Julia uses the first session category as the reference category. Thus in these tables “dms” can be interpreted as “session early & dms”.

In tables 3.1 to 3.6 the variable dms is the mean-subtracted session 0 DMS contrast dependent stimulus response magnitude. (L2-norm of the difference of the 100% contralateral stimulus contrast response to the 6%). session & dms denotes the interaction of the variables session and dms. The dependent variables: β_*contra*_, β_*ipsi*_, β_*bias*_ correspond to the behavioral model choice weights.

In tables 4.1 to 4.2 the variable cohort denotes the group identity of each mouse, either Chrmine or YFP. The variable contra denotes whether the trial corresponded to a stimulus contralateral from the recording site. The dependent variable *correct* is the side-specific (contra or ipsi) accuracy. Interactions and reference levels are as described above, thus the term: cohort: chrmine & contra:ipsi is the 3-way interaction of the reference level for session (sessions 1-7, e.g. “session early”), cohort, and contra.

## Supplemental Figures

**Supplemental Figure 1.**
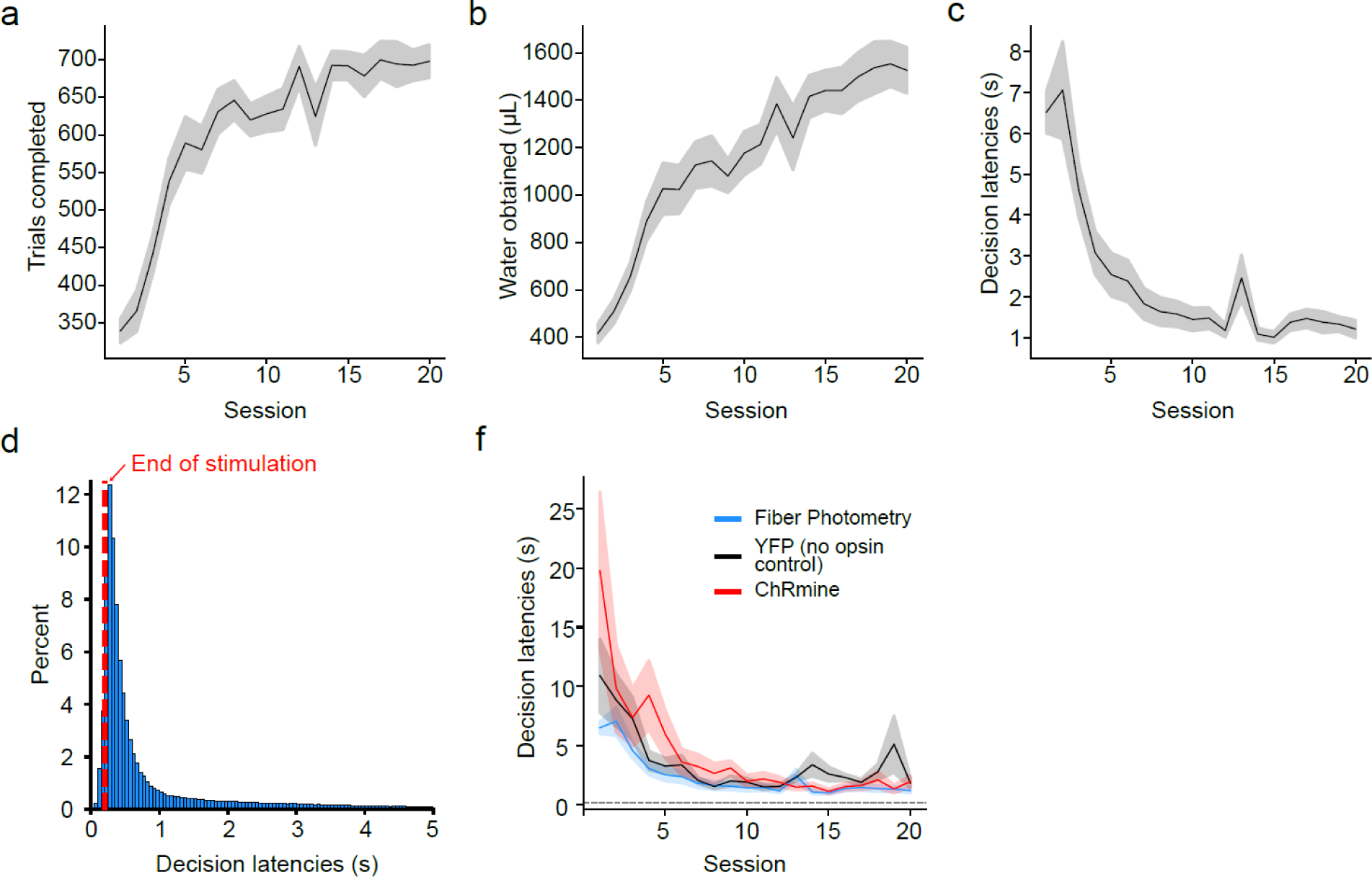
Session statistics throughout training, including sessions with optogenetic stimulation (related to figure 1 and 4). **a.** Trials completed, **b.** Total water obtained across sessions. **c.** Decision latencies (time between the go cue and the outcome delivery) across sessions. **d.** Histogram of decision latencies across all mice in Fig. 1-3 (*n=22*). Red line denotes the end of the optogenetic stimulation trains (0.2 s from the go cue) in Fig. 4. **e.** Comparison of decision latencies across sessions across ChRmine stimulation (red, *n=7*, Fig. 4), no opsin control (YFP, black, *n=6*, Fig. 4) and the fiber photometry (blue, *n*=*22*, Fig. 1-3) cohorts. Dashed line represents the end of the optogenetic stimulation (200 ms). Across all panels, lines and shading represent mean +/- s.e.m across mice.

**Supplemental Figure 2.**
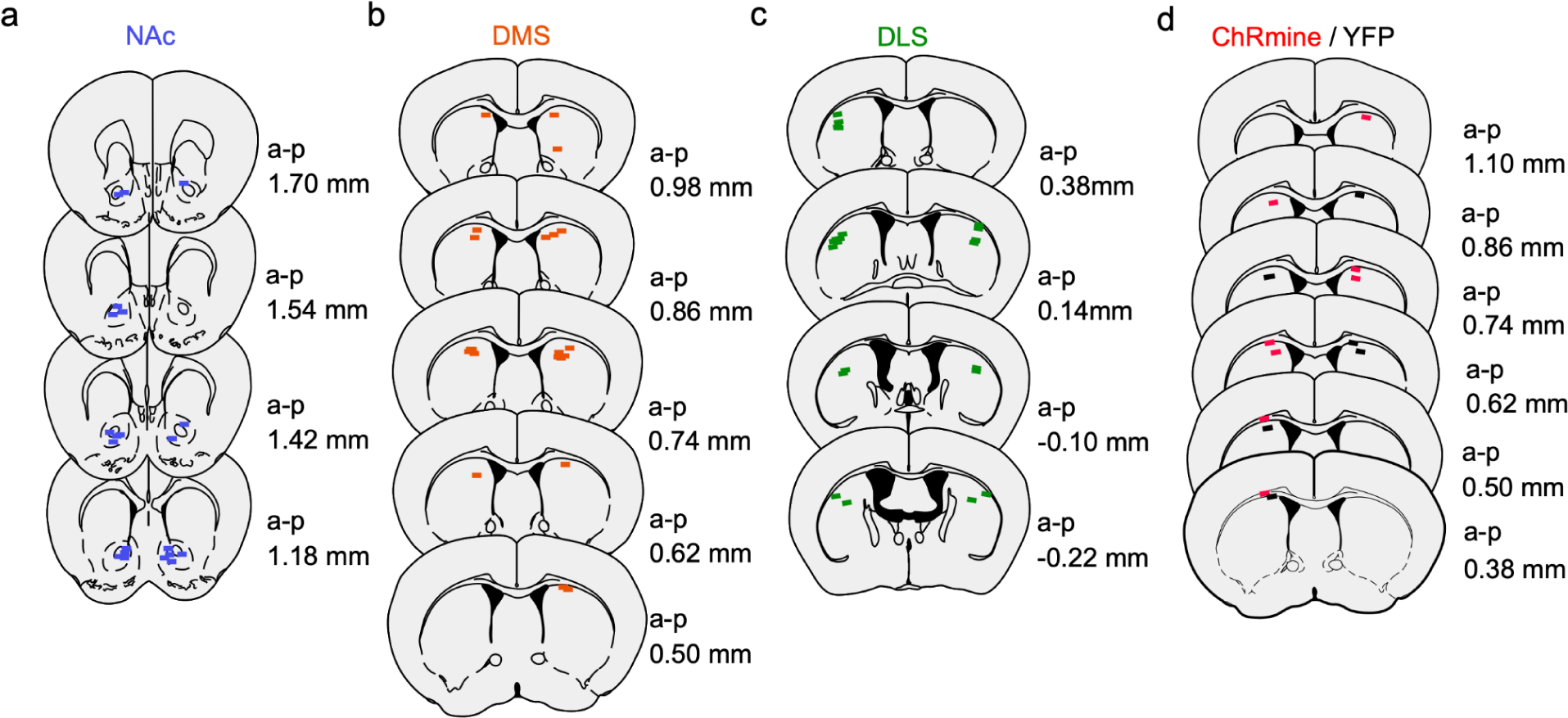
Optical fiber location for dopaminergic terminal recordings and terminal optogenetic stimulation (related to Figure 2 - 4). Received fiber tip locations for the fiber photometry recordings in Fig. 2-3 in **a.**, NAc, **b.**, DMS and **c.**, DLS. Each line (200µm) represents a fiber tip and their color relays their assigned striatal subregion: Blue - NAc, Orange -DMS, Green - DLS. **d.** Recovered fiber tip locations for the optogenetic terminal stimulation experiment in Fig. 4. Each line (300µm) represents a reconstructed fiber tip and their color relays their assigned group: Black - YFP (no opsin control) or Red - ChRmine. All fibers were located to the closest 100 µm section in the Paxinos-Franklin atlas ^94^. Sections are ordered by anterior-posterior (a-p) distance from Bregma.

**Supplemental Figure 3.**
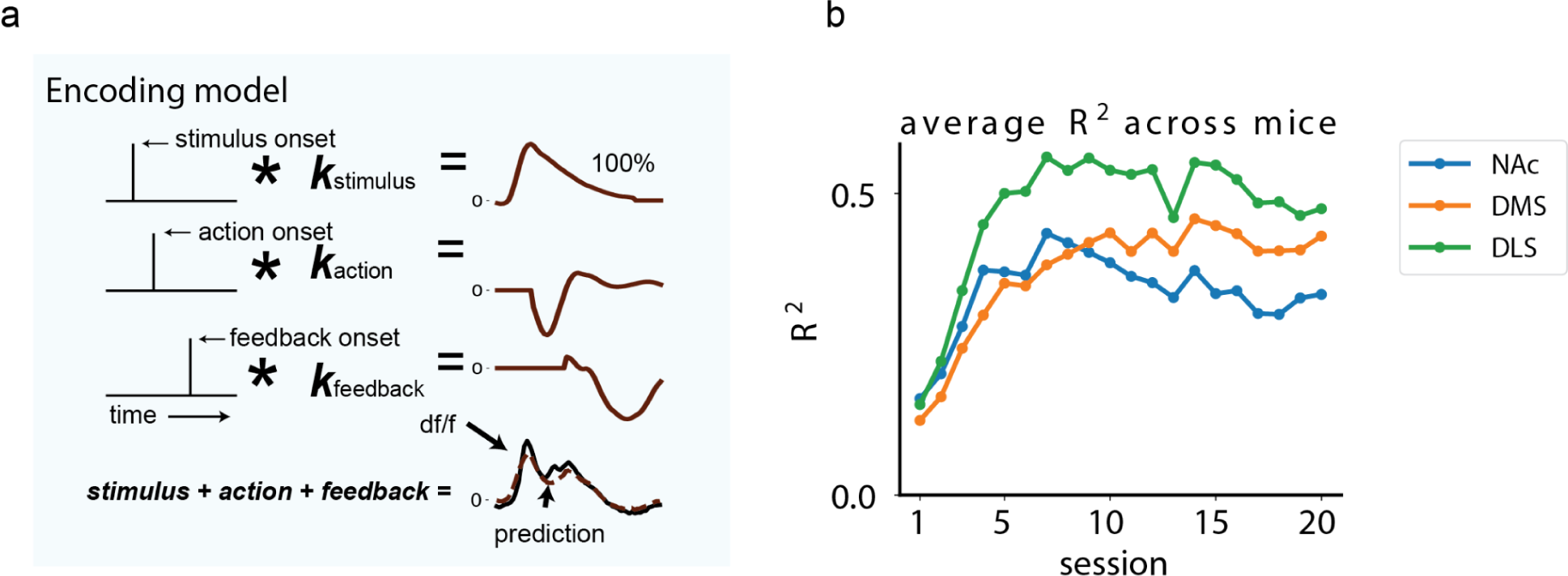
Encoding model schematic & average explained variance (related to Figure 2) **a.** Encoding model schematic (see Methods for details). We convolved delta functions defining task relevant events such as stimulus onset, action onset, and feedback onset with temporal kernels of those events, then summed up all components to get the predicted response. Example 100% contrast trial is shown. **b.** Explained variance in the fluorescence data (dF/F) by the model predictions, averaged across mice per training session. *R*^2^ is the variance explained across all trials within a session (from stimulus onset to 1 second after feedback for each trial).

**Supplemental Figure 4.**
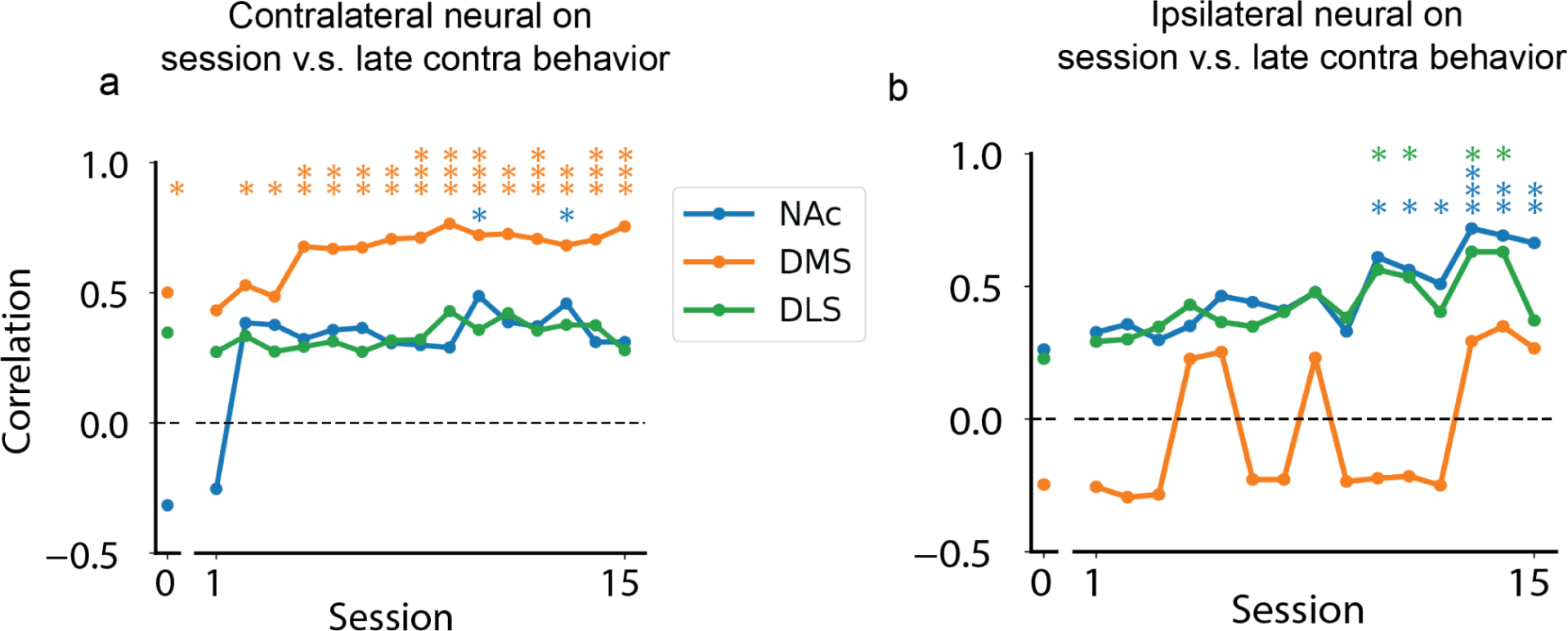
Throughout training, contrast-dependent contralateral DMS DA stimulus responses predict later contralateral stimulus-dependent behavior (related to Figure 3). **a.** For each region, and for each session, correlation across animals of the average contralateral stimulus sensitivity weight from the behavioral model at the end of training (sessions 16-20) with the contrast-dependence of the contralateral stimulus kernel (difference of L2-norm of the highest and lowest contrast contralateral stimulus response). Session 0 denotes the pre-exposure session before the start of training, described in Fig. 3. **b.** Same as a, however for ipsilateral (rather than contralateral) neural and behavioral weight estimates. In both panels, correlations and p-values are computed with robust regression, as described in the Statistical Analysis section of Methods.

## Acknowledgements

We thank Oliver Huang, Angela Chan, and the PNI Viral Core Facility for AAV production; Adrian Sirko and the Princeton Laboratory Animal Resources staff for help with animal husbandry; Zoe Ashwood, Sam Zorowitz, and Harrison Ritz for helpful discussion with behavioral modeling; and V. Roser, M. Siniscalchi, R. Fetcho and other members of the Witten lab for feedback on this work. This work was supported by NIH 1DP1MH136573 (IBW), the Simons Collaboration on the Global Brain (IBW, JP), Princeton Innovation Fund (IBW, ND), Wellcome Trust - Sir Henry Wellcome Fellowship 221657/Z/20/Z (AP-V), and 1U19NS123716-01 (IBW, JP).

